# Interactions in milk suggest a physiological role for β-lactoglobulin

**DOI:** 10.1101/675587

**Authors:** J.M. Crowther, M. Broadhurst, T. Laue, G.B. Jameson, A.J. Hodgkinson, R.C.J. Dobson

**Affiliations:** University of Edinburgh; AgResearch; University of New Hampshire; Massey University; University of Canterbury

## Abstract

β-Lactoglobulin is the most abundant protein in the whey fraction of ruminant milks, yet is absent in human milk. It has been studied intensively due to its impact on the processing and allergenic properties of ruminant milk products. However, the physiological function of β-lactoglobulin remains unclear. Sedimentation velocity experiments have identified new interactions between fluorescently-labelled β-lactoglobulin and other components in milk. Co-elution experiments support that these β-lactoglobulin interactions occur naturally in milk and provide evidence that the interacting partners are immunoglobulins, while further sedimentation velocity experiments confirm that an interaction occurs between these molecules. Ruminants (e.g. cows and goats) are born without circulating immunoglobulins, which they must obtain from their mothers’ milk, whilst humans obtain immunoglobulins both through milk and during gestation via the placenta. We propose that β-lactoglobulin serves to protect immunoglobulins within ruminant milk during digestion, ensuring their efficient transfer from mother to offspring.

**Statement of Significance:** β-Lactoglobulin is an abundant protein in the whey fraction of ruminant milks (e.g. cow and goat milk), yet it is completely absent in human milk. While this protein has been extensively studied, due to its impact on the processing and allergenic properties of milk, its physiological function remains unclear. We fluorescently labelled β-lactoglobulin to monitor its interactions with other milk components within its physiological environment, milk. Under these near physiological conditions β-lactoglobulin is capable of interacting with several classes of immunoglobulins. Immunoglobulins are susceptible to digestion, but are required to confer immunity from the mother to the offspring. We propose that β-lactoglobulin serves to protect immunoglobulins within ruminant milk during digestion, ensuring their efficient transfer from mother to offspring.

## Introduction

β-Lactoglobulin is the most abundant protein in the whey fraction of ruminant milk, including cow and goat milk, yet is absent in human milk. β-Lactoglobulin is known to affect the processing of ruminant milk, for instance due to heat-induced aggregation during heat treatment (1). It is also one of the main immunogenic proteins that contribute to milk allergies (2). Considerable effort has been made to elucidate the biological and biophysical properties of β-lactoglobulin and as such it is one of the most highly studied proteins, featured in the title or abstract of over 4000 publications in the last 75 years. Despite many conjectures, the physiological role of β-lactoglobulin remains a mystery.

β-Lactoglobulin belongs to the lipocalin family, which are a group of small extracellular proteins. Despite considerable diversity at the sequence level, lipocalin proteins share a conserved protein fold: an eight stranded β-barrel enclosing a large hydrophobic cup-shaped cavity (termed a calyx), together with a 3-turn helix between the seventh and eighth β-strands (3). This fold makes lipocalins well suited to binding a range of hydrophobic molecules. While once simply classified as transport proteins, it is now known that lipocalins exhibit great functional diversity (3–10).

β-Lactoglobulin was first thought to function as a transporter of retinol between mother and offspring, after it was demonstrated that bovine β-lactoglobulin can bind retinol (vitamin A) (11). Since then, bovine β-lactoglobulin has been shown to bind a range of small hydrophobic molecules, including vitamins, cholesterol and a range of fatty acids, as reviewed by Le Maux et al. (12) and Kontopidis et al. (13). The apparent lack of selectivity makes it less likely that β-lactoglobulin is a specific fatty acid or vitamin transporter. Further, given the absence of β-lactoglobulin in the milk of humans, it is unlikely that this protein participates in a process that is still required in humans.

The closest human homologue to β-lactoglobulin is glycodelin (also known as pregnancy protein 14) (14). Unlike β-lactoglobulin and other lipocalins, glycodelin is a glycoprotein. Glycosylation is crucial for the various functions of glycodelin, which include an immunosuppressive activity in the uterus to protect products of the reproductive organs from the immune system. The lack of glycosylation makes it unlikely that β-lactoglobulin is capable of fulfilling a similar role.

β-Lactoglobulin may simply act as a source of amino acids for the offspring of the animals that produce it. However, the resistance of β-lactoglobulin to digestion by human proteolytic enzymes (15) and the presence of highly conserved features, including a pH-gated loop movement, among β-lactoglobulin orthologues (16) argue against a simple nutritive function.

There have been conflicting reports on the antimicrobial activity of β-lactoglobulin. Chaneton et al. (17) observed that isolated β-lactoglobulin inhibited the growth of *Staphylococcus aureus.* Peptides resulting from the enzymatic digestion of β-lactoglobulin appear to possess some antibacterial activity against both *Escherichia coli* and *S. aureus in vitro* (18,19). However, while Fijalkowski et al. (20) observed that incubation of *S. aureus* in whey resulted in growth inhibition, they found no substantial effect using pure β-lactoglobulin.

We reasoned that the native function of β-lactoglobulin might be uncovered if its physiological interactions were known. The use of fluorescence detection analytical ultracentrifugation allowed us to monitor the interactions of β-lactoglobulin in milk, a highly complex colloid, for the first time. The sedimentation of fluorescently-labelled bovine and caprine β-lactoglobulin are both significantly altered in the presence of cow and goat milk, a signature that they interact with other components. Co-elution experiments with milk further support the presence of these interactions. The results lead us to propose a new physiological role for β-lactoglobulin that explains the absence of this otherwise abundant protein from human milk.

## Methods

### Protein purification

The cloning, expression and purification of recombinant bovine β-lactoglobulin A and caprine β-lactoglobulin have been described in detail previously (21,22).

### Fluorescent labelling of proteins

Proteins were labelled with fluorescein isothiocyanate (FITC) by reacting proteins at a concentration of 5-10 mg/mL (in 0.1 M sodium carbonate buffer, pH 9) with 100 µL FITC per 1 mL of sample. FITC was dissolved in dimethylformamide at 10 mg/mL immediately prior to use. The solution was rotated to mix for one hour at ambient temperature in the dark, then passed through a 5 mL desalting column (GE Healthcare) to remove the bulk of the free label. β-Lactoglobulin was further purified by gel filtration chromatography (HiLoad Superdex 200 16/60 120 mL). The degree of labelling was calculated by measuring the absorbance at 280 nm and 494 nm using a NanoDrop™ spectrophotometer. The protein concentration was calculated from the absorbance at 280 nm taking into account the contribution of FITC, as per Eq. 1.

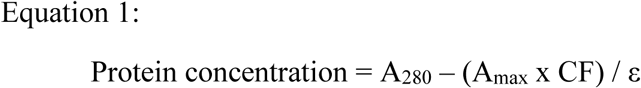

Where A_max_ is the wavelength of maximum absorbance for the dye molecule (494 nm for FITC), CF is the correction factor which adjusts for the amount of absorbance at 280 nm caused by the dye (0.3 for FITC, as supplied by Thermo Fisher Scientific), and ε is the protein molar extinction coefficient (17210 M^-1^ cm^-1^ for both bovine and caprine β-lactoglobulin, as calculated using ExPasy ProtParam (23)).

The degree of labelling was calculated as according to Eq. 2. A degree of labelling between 0.3 and 1.2 moles of dye per mole of protein was deemed acceptable.

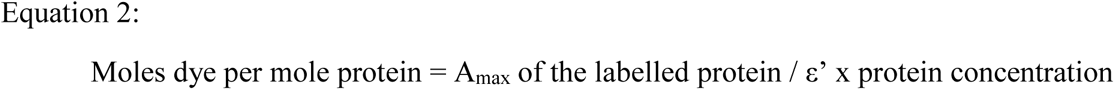

Where ε’ is the molar extinction coefficient of the fluorescent dye (68000 M^-1^ cm^-1^ for FITC, as supplied by Thermo Fisher Scientific).

### Analytical ultracentrifugation

Sedimentation velocity experiments were conducted in a Beckman Coulter XL-I analytical ultracentrifuge. Depending on the experiment, sedimentation was monitored utilising one of the three available optical systems (absorbance, interference and fluorescence). Specific experimental conditions, i.e. the buffer used, the protein concentrations, and the wavelengths measured, are specified in each of the figure legends. Reference solution (400 µL) and sample solutions (380 µL) were added to 12 mm double sector cells with quartz or sapphire windows. Cells were mounted in an An-50 Titanium eight-hole rotor. Initial scans were performed at 3,000 rpm to determine the optimal settings, with sedimentation performed at 50,000 rpm at 20 °C. Data were collected in step sizes of 0.003 cm with no delay between scans and no averaging.

Buffer density and viscosity and an estimate of the partial specific volume of proteins were calculated using SEDNTERP (24). Data were fitted to a continuous sedimentation coefficient [*c*(s)] model or continuous mass [*c*(M)] model using SEDFIT (25). Data were also subjected to two-dimensional spectrum analyses with genetic algorithm optimisation, and van-Holde/Weischet analyses in UltraScan III (26–28).

### Preparation of milk for interaction studies

Fresh, raw, cow milk (Holstein breed) was obtained from the Fairchild Dairy Teaching and Research Centre at the University of New Hampshire, United States of America. Fresh, raw, goat milk was obtained from a local farmer in either New Hampshire, United States of America (Nubian breed) or Canterbury, New Zealand (Nubian/Saanen cross breed). Before use, the milk was spun at 5000 g for 5 mins. The layer of fat that resulted at the top was removed and the liquid underneath was transferred to a new vessel, leaving behind the small pellet of cells that formed at the bottom of the milk sample. Diluted skim milk samples were used for all AUC studies, using 0.1 M sodium phosphate, 0.1 M NaCl, pH 6.7 buffer.

### Size-exclusion chromatography

Samples of skim cow milk were subjected to size-exclusion chromatography using a Superdex 200 PG 1800 column, with 15 mL fractions collected. A dotblot grid (1 × 1 cm) was marked onto a nitrocellulose membrane (Biotrace NT). 2 μL of each eluted fraction was spotted per square, with 2 μL of diluted loading sample loaded as a comparison. The membrane was allowed to air dry, then peroxidase activity was blocked with 3% hydrogen peroxide/1% sodium azide solution at room temperature for 10 minutes. After washing, the membrane was blocked with either 1% polyvinylpyrrolidone-25 (PVP-25) or 1% bovine serum albumin (BSA) in Tris-buffered saline, pH 7.6, containing 0.1% Tween-20 and 0.1% BSA, at room temperature for 2-3 hours. The membrane was incubated with anti-bovine β-lactoglobulin horse radish peroxidase (HRP)-conjugated antibody (Bethyl Lab A10-125P 1:100K) for 3 hours and immune-reactivity determined using chemiluminescence. In a similar manner, reactivity of eluted fractions was determined versus anti-bovine IgM (Bethyl A10-101P 1:40K), anti-bovine IgG (Bethyl Lab A10-118P 1:75K) and anti-bovine IgA (Bethyl Lab A10-131P 1: 40K) antibodies.

## Results

### Characterising the behaviour of β-lactoglobulin in cow and goat milk

To identify interactions between β-lactoglobulin and other components in milk, recombinantly expressed and purified bovine and caprine β-lactoglobulin were labelled with fluorescein isothiocyanate (FITC), before being added to samples of diluted cow and goat milk. The sedimentation of β-lactoglobulin, monitored using the fluorescence-optics within the analytical ultracentrifuge, is significantly faster in the presence of cow or goat milk when compared to equivalent studies in simple buffered solutions. This suggests that β-lactoglobulin is forming higher molecular weight species with other milk components.

A dilution series of cow and goat milk samples was prepared, to which fluorescently-labelled bovine or caprine β-lactoglobulin was added to a final concentration of 0.75 µM. This is much lower than the concentration of β-lactoglobulin usually found in milk (∼150 µM (12)). At this concentration (0.75 μM) and pH (6.7), bovine and caprine β-lactoglobulin are predominantly monomeric with a small amount of dimer present (see the black lines in Fig. 1). The sedimentation profile changes considerably once milk is added to the solution, with the appearance of a new, larger species with a sedimentation coefficient of ∼8-10 S (Fig. 1). As the concentration of milk increases a greater proportion of labelled β-lactoglobulin is found in complex with other components. The sedimentation velocity data were also analysed using the van-Holde/Weischet method: a model-independent analysis that directly assesses the shape of the sedimenting boundary. These analyses support the conclusion made from the *c*(s) analyses that with increasing concentrations of milk the proportion of β-lactoglobulin that is found in complex increases (Fig. 1 E-H).

**Figure 1:**
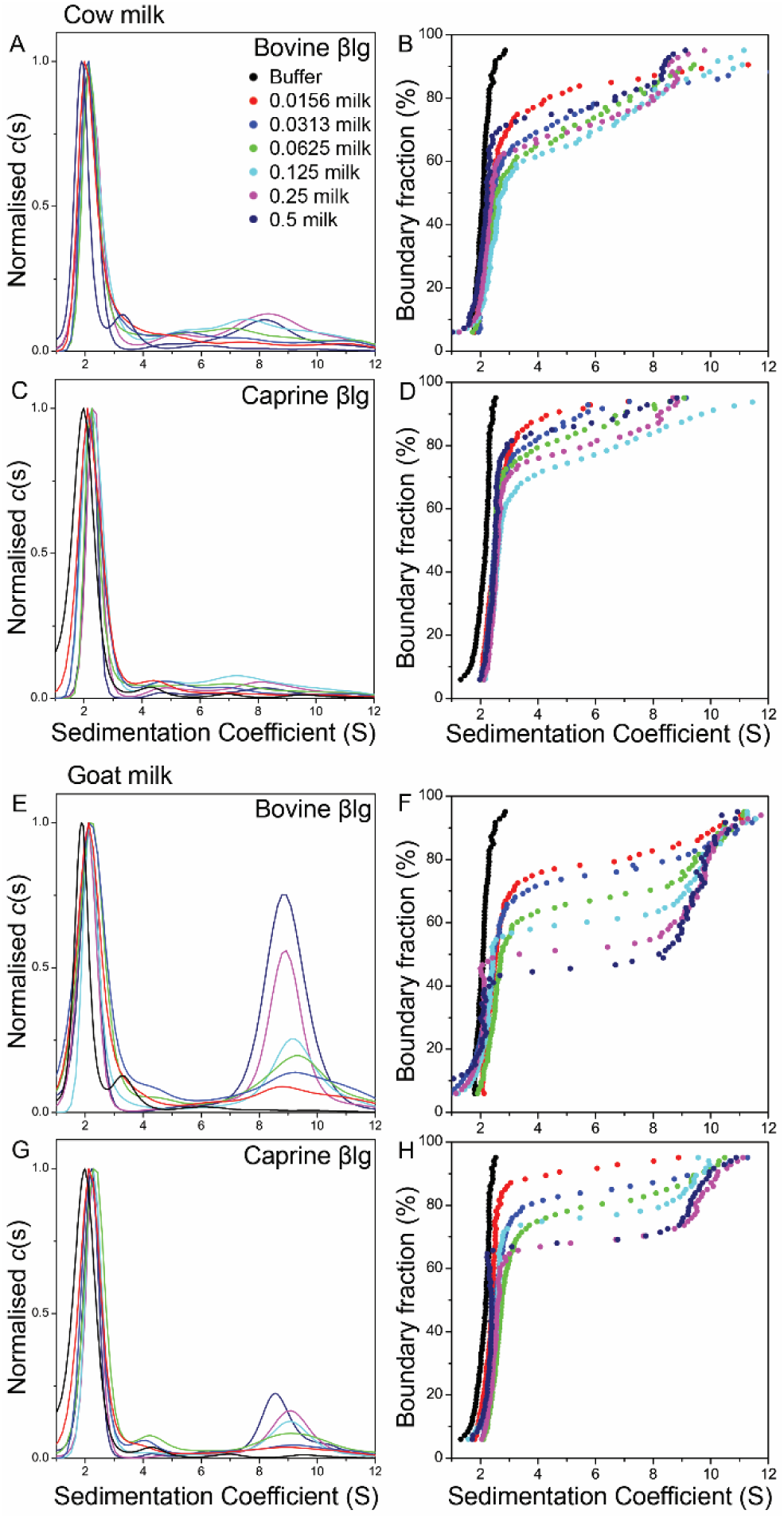
Sedimentation velocity analysis of bovine and caprine β-lactoglobulin in cow and goat milk. Each milk was serially diluted from ½ (0.5) to 1/64 (0.0156) dilution. FITC-labelled protein was added to each sample to a final concentration of 0.75 µM. The data were fitted to a continuous sedimentation coefficient distribution (*c*(*s*)) model for A) FITC-labelled bovine β-lactoglobulin A in cow milk, B) FITC-labelled caprine β-lactoglobulin in cow milk, C) FITC-labelled bovine β-lactoglobulin A in goat milk, D) FITC-labelled caprine β-lactoglobulin in goat milk. E-H) the same data were analysed using the van-Holde/Weischet method.

Different behaviours were observed for both the type of milk used and the orthologue of β-lactoglobulin examined. In each milk a greater amount of complex is formed for bovine β-lactoglobulin than for caprine β-lactoglobulin (compare Fig. 1A and 1C to Fig. 1B and 1D), despite the proteins having a very high level of sequence and structural identity (22). As the same concentration of β-lactoglobulin was added to each sample, this suggests that the A isoform of bovine β-lactoglobulin (used in these interaction studies) has a higher binding affinity for the interacting component than caprine β-lactoglobulin. In goat milk, a greater amount of complex is formed with both bovine and caprine β-lactoglobulin as compared to that seen in cow milk (compare Fig. 1C and 1D to Fig. 1A and 1B). As the same volume of milk was added to each sample, this suggests that the interacting component has a higher affinity for β-lactoglobulin or is present in goat milk at a higher concentration.

The sedimentation of free FITC dye is not altered in the presence of milk (Fig. 2A). This demonstrates that FITC is not interacting with other components in milk, and is thus not responsible for the peak seen at 8-10 S in Fig. 1. We expect that fluorescently-labelled β-lactoglobulin is capable of mixing with endogenous β-lactoglobulin already present in milk, forming hetero-dimers, which would effectively increase the concentration of fluorescently-labelled β-lactoglobulin. Therefore, it is possible that the larger species formed in the presence of milk is a product of the self-association of β-lactoglobulin into an even higher-order oligomer. However, sedimentation velocity experiments of bovine β-lactoglobulin and caprine β-lactoglobulin at high concentrations (up to 400 µM) in buffered solutions unequivocally show a single species with a sedimentation coefficient of 2.6 S (Fig. 2B). This demonstrates that β-lactoglobulin does not form any higher-order species above a dimer, even at the concentrations that may be encountered in milk.

**Figure 2:**
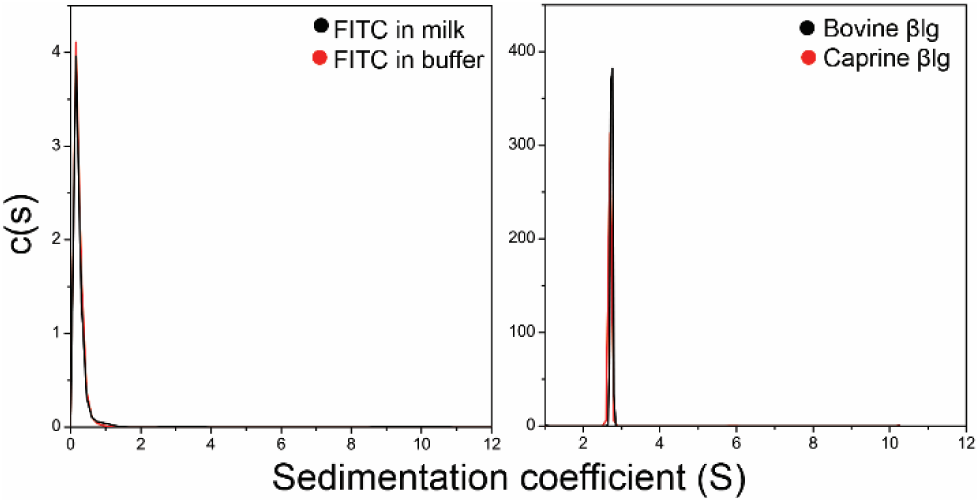
A) Sedimentation velocity analysis (using absorbance optics) of FITC in buffer and in goat milk (1/20 dilution). B) Sedimentation velocity analysis (using interference optics) of bovine and caprine β-lactoglobulin at 400 µM in buffer (0.1 M sodium phosphate, 0.1 M NaCl, pH 6.7).

Unlabelled β-lactoglobulin, added to milk that already contains FITC-labelled β-lactoglobulin, can compete for the interacting site that leads to the higher molecular mass species. This can be seen in Fig. 3 as a reduction in peak size at 8 S (Fig. 3A and 3B) and a decrease in the fraction of the boundary corresponding to the larger species (Fig. 3C and 3D). This demonstrates that the formation of the higher molecular mass species at 8 S is not due to any structural changes in β-lactoglobulin induced by the addition of FITC. Curiously, the sedimentation coefficient of the larger species appears to decrease, with increasing concentration of unlabelled β-lactoglobulin (Fig. 3), suggesting that the complex is decreasing in size. This could be attributed to a higher viscosity and density of the solution as the concentration of unlabelled protein is increased. Alternatively, it may suggest a network is formed between β-lactoglobulin and the interacting partner (which would imply multiple binding sites on both β-lactoglobulin and the interacting partner), where excess of either partner would disrupt the network and result in a reduced average complex size.

**Figure 3:**
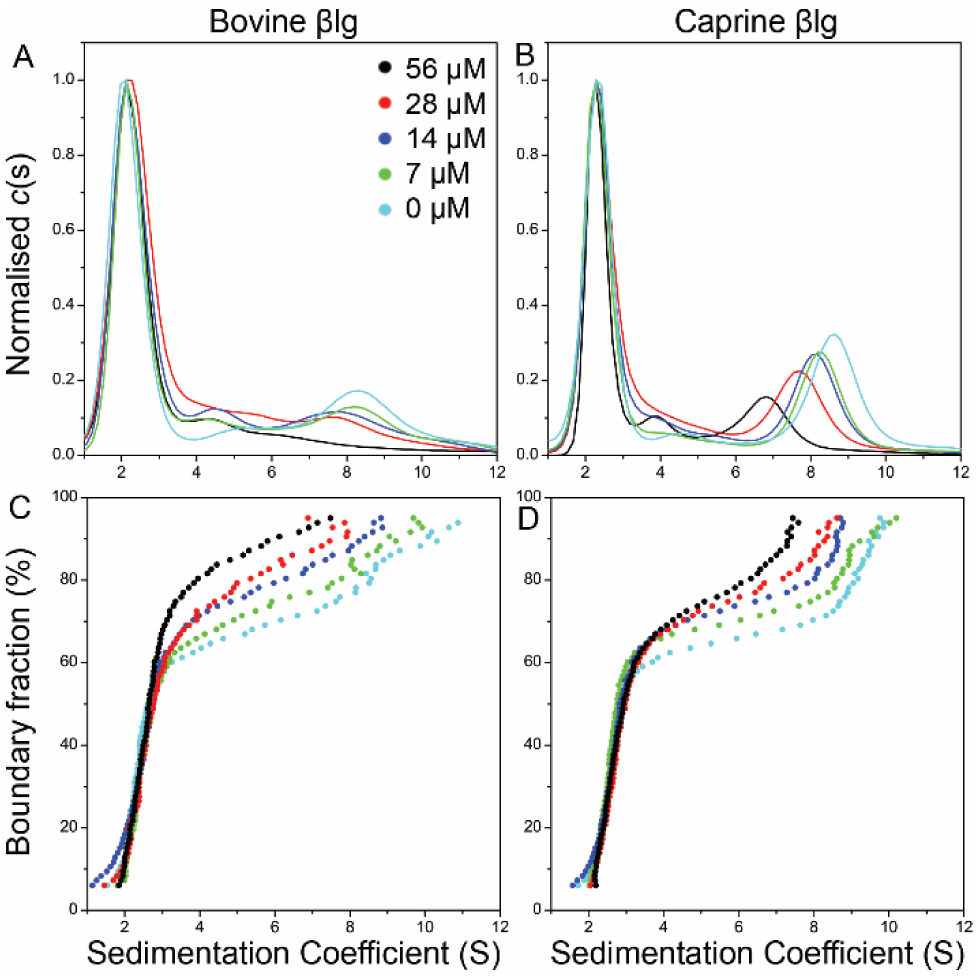
Sedimentation velocity analysis of FITC-labelled β-lactoglobulin proteins (0.75 µM) in milk (1/4 dilution) with increasing concentrations of unlabelled β-lactoglobulin (0–56 µM). Continuous sedimentation coefficient distributions of A) FITC-labelled bovine β-lactoglobulin A in cow milk with unlabelled bovine β-lactoglobulin A, and B) FITC-labelled caprine β-lactoglobulin in goat milk with unlabelled caprine β-lactoglobulin. C and D) enhanced van-Holde/Weischet analyses.

The higher molecular mass species associated with β-lactoglobulin can also be observed in an unmodified sample of milk when analysed by gel-filtration chromatography. β-Lactoglobulin was identified in the eluted fractions by means of a Western dot-blot assay using antibodies specific for bovine β-lactoglobulin. It is apparent that β-lactoglobulin elutes in two places consistent with our sedimentation experiments: late in the elution consistent with free β-lactoglobulin (see Box A in Fig. 4A), and earlier in the elution at a location expected for a much higher molecular mass species (Box B, Fig. 4A). This suggests that β-lactoglobulin is present in milk as two populations of different sized species, which agrees directly with what is seen in the AUC experiments. The importance of this result lies in the fact that the size-exclusion chromatography separation involves endogenous β-lactoglobulin that has not been recombinantly expressed or modified in any way, yet the same outcome is seen.

**Figure 4:**
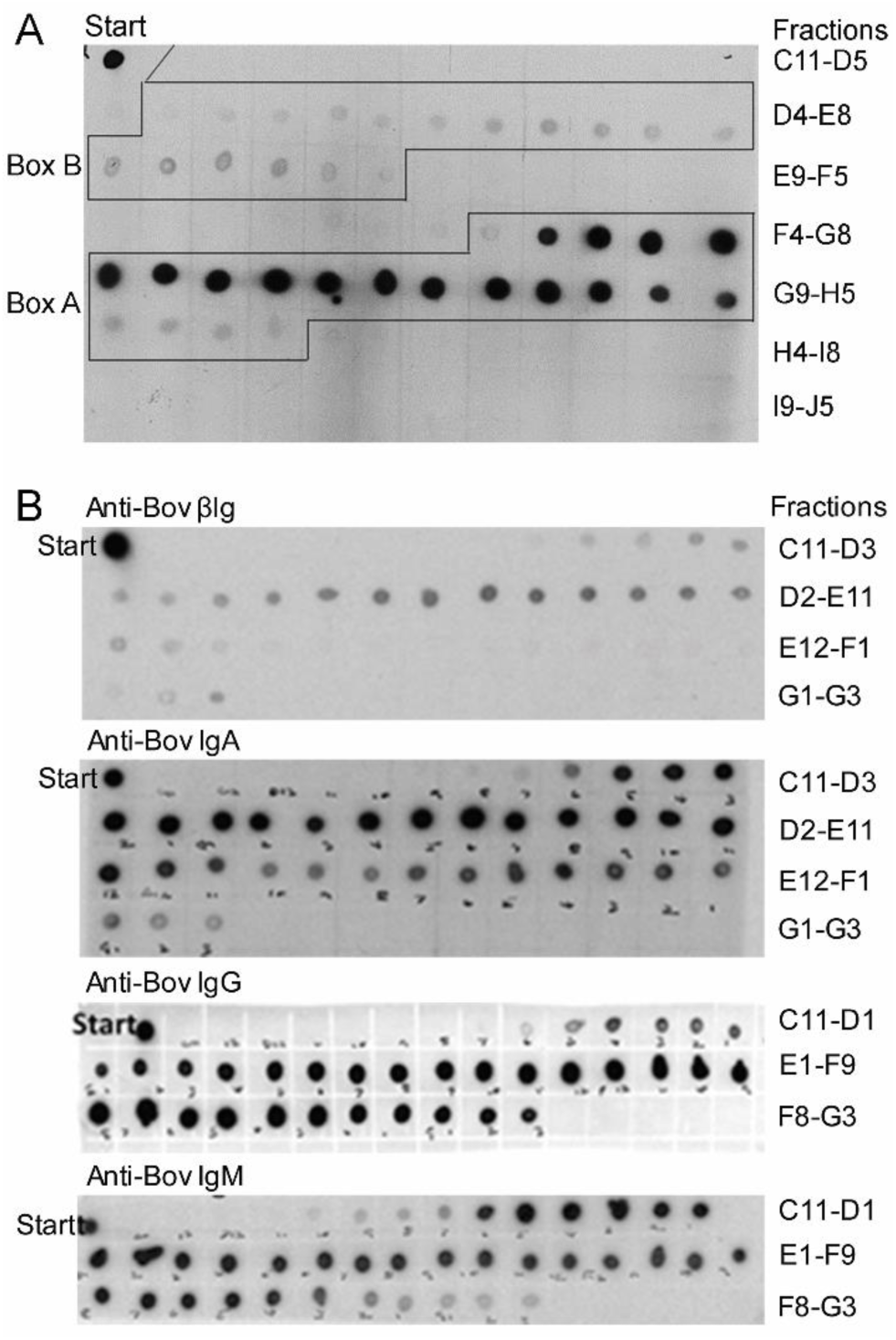
A) Western dot-blot of cow milk fractions following size-exclusion chromatography, utilising anti-bovine β-lactoglobulin antibodies. B) Western dot-blot of the same milk fractions utilising anti-bovine β-lactoglobulin, anti-bovine IgA, anti-bovine IgG and anti-bovine IgM.

In summary, the formation of a higher molecular mass species has been observed between bovine and caprine β-lactoglobulin, and other components within cow and goat milk. The possibility of self-association of β-lactoglobulin into a higher order species is ruled out. FITC molecules attached to the surface of β-lactoglobulin proteins are not responsible for complex formation, demonstrated by the fact that FITC does not interact with any milk components and that unlabelled β-lactoglobulin proteins can successfully compete for binding with labelled β-lactoglobulin. Even more convincingly, a higher molecular mass species of β-lactoglobulin can be seen in milk that contains β-lactoglobulin that has not been altered in any way.

### Identifying the interacting component

While bovine and caprine β-lactoglobulin can both form associations with kappa-casein in milk, this interaction is heat-induced and involves a disulfide-exchange reaction (29). Thus, the 8 S species identified in our sedimentation velocity and size-exclusion experiments is unlikely to be an association between β-lactoglobulin and kappa-casein. Given the size of the complex (8-10 S), we hypothesised that β-lactoglobulin interacts with immunoglobulin proteins present in milk.

The fractions of milk eluted from size exclusion chromatography, as described above, were also probed with anti-bovine IgG, IgA and IgM antibodies. It is apparent that these immunoglobulins co-elute from milk in the same fractions as the higher molecular mass bovine β-lactoglobulin (Fig. 4B). This is consistent with bovine β-lactoglobulin binding immunoglobulins in milk to form the higher molecular mass species seen in the sedimentation velocity experiments.

To confirm that β-lactoglobulin interacts with immunoglobulins, the sedimentation of bovine and caprine β-lactoglobulin was analysed in the presence of bovine and caprine IgG, purified from the serum of non-immunised cows and goats. For this experiment the sedimentation of FITC-labelled β-lactoglobulin was monitored using absorbance at 495 nm, using the AUC absorbance optical system. It can be seen that the sedimentation of both FITC-labelled bovine and caprine β-lactoglobulin is indeed altered in the presence of IgG molecules (Fig. 5), with a larger species appearing in the *c*(*s*) distribution at 6 S. While this is not as large as the 8-10 S species observed in the milk interaction experiments, this is likely due to these experiments being carried out on purified proteins in a simple buffered solution rather than within the complex milieu of milk.

**Figure 5:**
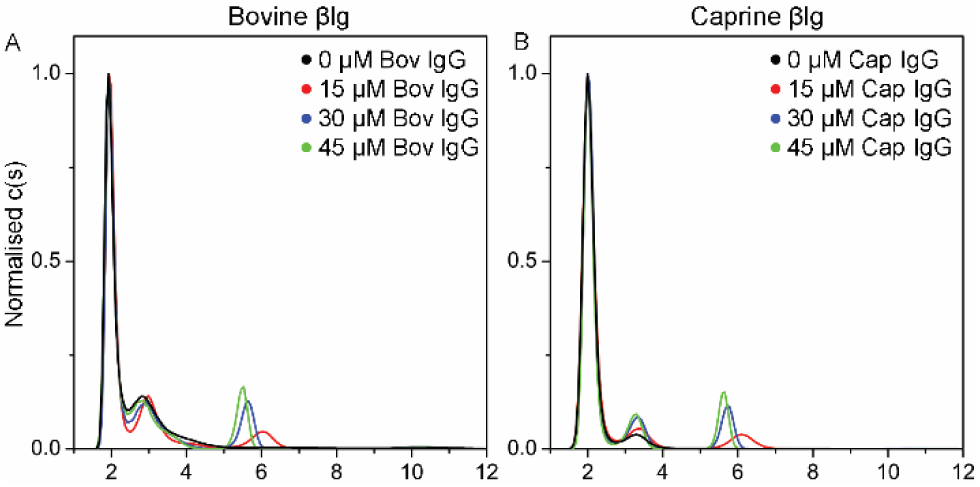
Sedimentation velocity analysis of A) FITC-labelled bovine β-lactoglobulin (10 µM) and bovine IgG (0-45 µM) and B) FITC-labelled caprine β-lactoglobulin (10 µM) and caprine IgG (0-45 µM).

To summarise, the co-elution of β-lactoglobulin with immunoglobulins IgG, IgA and IgM from cow milk following size-exclusion chromatography supports the hypothesis that β-lactoglobulin interacts with immunoglobulins within milk. The sedimentation of β-lactoglobulin is altered in the presence of IgG, strong evidence of an interaction between these two species.

## Discussion

We propose that the physiological function of β-lactoglobulin within ruminant milks is to protect immunoglobulins, particularly IgG, from digestive enzymes as they traverse the digestive tract, leaving them available to provide immune protection to the ruminant offspring.

Given the abundance of β-lactoglobulin in ruminant milk, and its absence in the milk of humans, it is unlikely that this protein would fulfil a role that is necessary in humans. In primates and lagomorphs (*e.g.* rodents and rabbits), species that lack β-lactoglobulin in their milk, IgG is transferred to the foetus *via* the placenta, and the offspring is born with circulating IgG (30). Conversely, the offspring of ruminants (such as cows and goats) are born agammaglobulinemic (*i.e.* with no circulating antibodies) and thus fully rely on the uptake of IgG from colostrum and milk to provide immune protection (30). Ruminant milk is therefore high in IgG (31).

We have shown here that bovine and caprine β-lactoglobulin are capable of interacting with immunoglobulins within cow and goat milk. Importantly, the resistance of β-lactoglobulin to gastric digestion is well known (15). The binding of β-lactoglobulin to immunoglobulins may serve to increase their resistance toward proteolysis during transit through the gastrointestinal tract. This would be particularly relevant for IgG, given its importance for transferring immunity in ruminants.

A structure of the complex between β-lactoglobulin and these immunoglobulins is necessary to fully understand the nature of this binding interaction. The crystal structure of bovine β-lactoglobulin in complex with an IgE Fab fragment has been reported (32). This complex demonstrates the interaction between an allergen and the antigen-binding region of the antibody, which provides structural insight into the recognition of this milk antigen by the human immune system. Immunoglobulins and β-lactoglobulin within milk would be expected to bind using a different mechanism. This could involve the heavy chain region of the immunoglobulin, rather than the Fab region, as the latter mechanism would likely elicit unwanted immune responses.

In conclusion, an interaction between β-lactoglobulin and immunoglobulins within both cow and goat milk has been identified for the first time, using analytical ultracentrifugation and size-exclusion chromatography experiments. We propose that this interaction with protease-resistant β-lactoglobulin protects immunoglobulins, essential to immunity of the neonate calf or kid, from proteolytic attack.

## Author contributions

Designed research: JMC, TL, GBJ, AJH, RCJD; performed research: JMC, MB; analyzed data: JMC, MB, TL, GBJ, AJH, RCJD; contributed analytic tools: TL; wrote the manuscript: JMC; edited the manuscript: TL, GBJ, AJH, RCJD.

## Acknowledgements

The authors declare no conflicts of interest.

